# Hypoactivation of ventromedial frontal cortex in major depressive disorder: an MEG study of the Reward Positivity

**DOI:** 10.1101/2024.04.18.590159

**Authors:** Christopher J.H. Pirrung, Garima Singh, Jeremy Hogeveen, Davin Quinn, James F. Cavanagh

## Abstract

**Background:** The Reward Positivity (RewP) is sensitive and specific electrophysiological marker of reward receipt. These characteristics make it a compelling candidate marker of dysfunctional reward processing in major depressive disorder. We previously proposed that the RewP is a nexus of multiple aspects of reward variance, and that a diminished RewP in depression might only reflect a deficit in some of this variance. Specifically, we predicted a diminished ventromedial contribution in depression in the context of maintained reward learning.

**Methods:** Here we collected magnetoencephalographic (MEG) recordings of reward receipt in 43 individuals with major depressive disorder (MDD group) and 38 healthy controls (CTL group). MEG allows effective source estimation due to the absence of volume conduction that compromises electroencephalographic recordings.

**Results:** The MEG RewP analogue was generated by a broad set of cortical areas, yet only right ventromedial and right ventral temporal areas were diminished in MDD. These areas correlated with a principal component of anhedonia derived from multiple questionnaires. Compellingly, BA25 was the frontal region with the largest representation in both of these effects.

**Conclusions:** These findings not only advance our understanding underlying the computation of the RewP, but they also dovetail with convergent findings from other types of functional source imaging in depression, as well as from deep brain stimulation treatments. Together, these discoveries suggest that the RewP may be a valuable marker for objective assessment of reward affect and its disruption in major depression.

## 1. Introduction

As one of two cardinal symptoms of major depression, anhedonia has long been recognized as a stable independent construct (1). Although traditionally interpreted as a loss of consummatory pleasure, recent approaches have re-interpreted anhedonia as an outcome of deficient liking, motivation, learning, or decision-making (2,3). These deficits can be considered sub-elements of a supraordinate deficit in reward processing (3), yet definitive parsing of these sub-types has proved elusive. Self-report and behavioral measures of anhedonia tend to capture this supraordinate level while not probing the specific elements affected (4–8). Biological markers of anhedonia may provide the ability to differentiate these symptom subtypes in depression (9).

One promising candidate biomarker is the Reward Positivity (RewP), an event-related potential component that is sensitively and specifically elicited by rewards. The RewP is enhanced by better-than-expected outcomes (10–12) as well as an individual’s subjective liking of the reward (13–17). These effective vs. affective influences on the RewP are conceptually in line with the anhedonic phenotypes of deficient learning vs. diminished liking. Indeed, the RewP is diminished in depressed individuals (18–22). Yet it remains unknown if a diminished RewP in depression reflects an impaired ability to learn, or if it indexes more hedonically-related affective deficits.

Given findings of intact learning ability in the context of a reduced RewP, we previously predicted that learning and liking contributed independent variance to the RewP, and that only the latter portion of variance may be affected by major depression (23). We further suggested that ventromedial prefrontal cortical (vmPFC) areas could be specifically implicated in this depression-related alteration (24). The source of the human RewP has not been rigorously tested, but it is also often assumed to be generated in the dorsal cingulate (25). While this may be partially true, the low frequency and large amplitude of this signal suggests a mechanism for combining inputs across widespread generative structures. Recent reports have suggested that perigenual and ventral midline cortices contribute meaningful variance to the RewP (23,26,27), but it is of course tenuous to infer any source separations from scalp EEG recordings.

In the present study, we used magnetoencephalography (MEG) to determine likely neural generators of the RewP. MEG is not affected by volume condition and thus affords more accurate source estimation. We further assessed the influence of anhedonia in Major Depressive Disorder (MDD) on these source generators, with a particular interest in ventral cortical areas.

Previous fMRI studies suggest a robust but complex representation of anhedonia in vmPFC. Some studies show hypoactivation to rewarding feedback (28–30) while the majority reveal a counterintuitive hyperactivation of these areas in association with anhedonia (29,31–40). In the current report, we demonstrate the first MEG evidence for distributed sources of the RewP. We further provide the first evidence that a reduced RewP in MDD may be specifically due to diminished ventral cortical activity. Furthermore, we provide evidence to motivate a novel testable hypothesis about the nature of countervailing vmPFC activation in MDD vs. anhedonia.

## 2. Methods and Materials

### 2.1 Participants

All participants were aged 18-55 years, were fluent in English, had no premorbid major medical or psychiatric conditions, no history of substance abuse, and were not currently taking medications. All procedures were approved by the Institutional Review Board at the University of New Mexico and all participants provided written informed consent. Participants were first screened by phone to determine potential eligibility. Then they completed a Structured Clinical Interview for DSM-5 (SCID), a battery of questionnaires on mood and anhedonia, and two tasks designed to measure anhedonic phenotypes: the Effort Expenditure for Reward Task (6) and the Probabilistic Reward Task ((7); hereafter we refer to this as the Pizzagalli task for differentiation from the MEG task described below). Participants who met SCID criteria for a major depressive disorder (MDD group: n=52, 35 female) or who did not meet criteria for any Axis I disorder (CTL group: n=38, 21 female) were invited to the MRI and MEG sessions described below. Due to scheduling difficulties and other data collection issues, some participants have partial datasets (Fig S1).

### 2.2 Procedure

In the MEG, participants performed a probabilistic learning task using different pseudo-randomly assigned character sets (Fig 1A). This task included a forced choice training phase followed by a subsequent testing phase (41). During the training phase the participants were presented with three stimulus pairs, where each stimulus was associated with a different probabilistic chance of receiving ‘Correct’ or ‘Incorrect’ feedback. These stimulus pairs (and their probabilities of reward) were termed A / B (80% / 20%), C / D (70% / 30%) and E / F (60% / 40%). All training trials began with a fixation cross displayed for 1000 ms. The stimuli then appeared for a maximum of 4000 ms, and disappeared immediately after the choice was made. A jittered inter-stimulus-interval between 300 and 500 ms followed. If the participant failed to make a choice within the 4000 ms, “No Response” was presented. Following a button press, either ‘Correct’ or ‘Incorrect’ feedback was presented for 1000 ms. MEG signals were locked to these feedback presentations. A Q-learning model was applied to these training phase data to examine differences in learning rates between groups (see (18) for details). A subsequent testing block was used to derive behavioral measures of reward vs. punishment learning. In the test phase, all possible stimulus pairs were presented eight times (120 trials total) with no feedback provided after selection. Reward seeking (“Go learning”) was defined as the accuracy of choosing A over C, D, E and F (i.e., seeking A), whereas punishment avoidance or “NoGo learning” was defined as the accuracy of choosing C, D, E and F over B (i.e., avoiding B).

**Figure 1.**
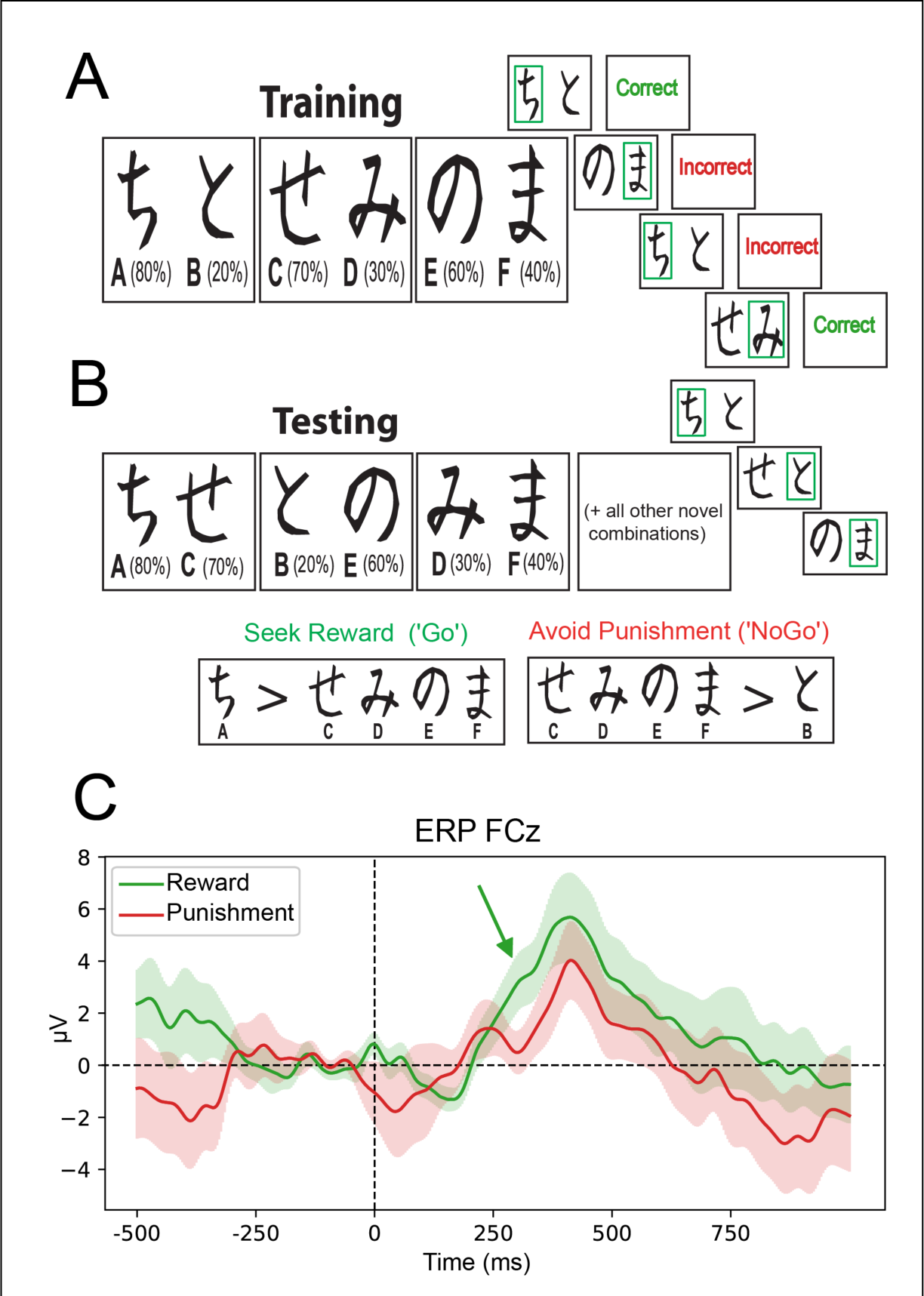
Probabilistic Selection Task design and ERPs. **A**) Training block task design with “Correct” and “Incorrect” feedback. **B)** Testing block task design without feedback. “Go” and “NoGo” learning represent reward seeking and punishment avoidance, respectively. **C)** Feedback locked ERPs at FCz following Reward and Punishment from participants with EEG data (N=24). The green arrow indicates the Reward Positivity.

### 2.3 MEG and EEG Acquisition and Sensor-Level Processing

MEG data were obtained using a 306-sensor Elekta Neuromag System. Data were recorded continuously at a sample rate of 1000 Hz across a frequency range of .1 to 330 Hz. In a subset of participants, EEG data were collected concurrently across a frequency range of .01 to 100 Hz using an EGI 128-electrode system with a sample rate of 250 Hz. EEG data were processed using EEGLab 2020 (42) in MATLAB R2022b (43). Data were re-referenced to a whole brain average, linearly detrended, then epoched around the presentation of a cue. Independent components analysis was used to remove blink artifacts from the data. Data were baseline corrected with a baseline window of 500 to 200 ms before cue, then time shifted to be feedback-locked. Epochs were then lowpass filtered at 20 Hz then averaged at FCz to create the ERP. Given the small subsample with concurrent EEG data, these data were only used to show that a RewP was elicited (Fig.1B).

MEG data were preprocessed using MNE-Python v.0.23.1 (44–46). Data were preprocessed using a Maxfilter and temporal signal-space separation (tSSS) to remove external noise and adjust for motion correction. Sensor-level data were also aligned to the device origin during this step. Of the 81 participants with usable MEG data for the PST, three participants had errors with this transformation and were excluded from sensor-level analyses (See Fig S1). Data were then high-pass filtered at 1 Hz to remove any slow drifts and notch filtered at 60 Hz to remove any powerline noise. Following this, data were down-sampled to 250 Hz. Signal-space projection (SSP) was then used to remove any artifacts from eye blinks and heart rate. Data were then epoched around the presentation of the cue.

Sensor-level MEG data were then baseline corrected to a period of 500 to 200 ms prior to cue then time shifted to be feedback-locked. Epochs were then lowpass filtered at 20 Hz and averaged to create sensor-level ERFs. The primary magnetometers for ERF presentation, MEG0511 and MEG0921, were chosen due to two factors: (1) magnetometers pick up deeper signals better than their hyper-focal gradiometer counterparts, (2) lateral sensors performed better than midline sensors at detecting medial signals, as the magnetic field is orthogonal to the direction of the electrical dipole. Sensor-level ERFs were used to determine the time window for source estimation, aided by computation of the global field power (GFP, see Fig S2).

### 2.4 MRI Acquisition and Processing

MRI images were collected using two scanners due to a scanner upgrade halfway through data collection. The first subset of scans consists of T1-weighted structural MRI images obtained with a Siemens 3T Trio TIM scanner and a 32-channel coil using a multi-echo MPRAGE sequence [TR/TE/TI = 2530/1.64, 3.5, 5.36, 7.22, 9.08/1200 ms, flip angle = 7°, field of view (FOV) = 256 × 256 mm, matrix = 256 × 256, 1 mm thick slice, 192 slices]. The second set of scans consisted of T1-weighted structural MRI images obtained with a Siemens 3T Magnetom PRISMA scanner and a 32-channel coil using a multi-echo MPRAGE sequence [TR/TE/TI = 2530/1.69, 3.55, 5.41, 7.27, 9.13/1200 ms]. Heudiconv was used to organize the data in BIDS format. Then data were processed with fMRIprep 1.5.0 software (47). MPRAGE sequences were first skull stripped, spatially normalized to MNI152NLin2009cAsym, tissue segmented, and then brain surface reconstruction was performed using Freesurfer (48).

### 2.5 MEG Source-Level Processing

Preprocessing and epoching for the MEG data were identical to sensor-level analyses, with the exception of aligning to device origin during tSSS. Source estimates were calculated using only magnetometers to bolster deeper source estimation. A surface source space for each individual was created from the T1 image using skull surfaces created by the Freesurfer watershed boundary element method (BEM) algorithm. This source space was used to create the forward solution. Source space was further limited to cortical areas by removing unlabeled areas on the MultiModal Parcellation atlas (HCP-MMP1; (49)). The epoched MEG data were then baseline corrected to a period of 500 to 200 ms prior to cue and low-pass filtered at 20 Hz. After creating a noise covariance matrix using this pre-cue baseline period, MEG data were time-shifted to the feedback. The inverse operator was then created using the forward solution, feedback-locked MEG data, and the source space. This inverse operator was then applied to each condition (reward/punishment) of the MEG data and projected onto an inflated brain, surface source space using dynamic statistical parametric mapping (dSPM). These projections were then morphed to the “fsaverage” brain for group display.

### 2.6 Source Statistics

Source estimates were regularized prior to group statistics by converting individual data to percent signal change. First, the dSPM contrast of reward > punishment was examined in all participants using a within-subject spatiotemporal permutation clustering test (permutations=5000, cluster threshold=0.001). This test was limited to 350-450 ms, as determined by global field power and the sensor-level ERFs. These clusters of significant vertices were then used as a mask, restricting the search space for an independent-sample permutation clustering test of the orthogonal CTL > MDD contrast. An additional spatiotemporal permutation clustering test (permutations=5000, cluster threshold=0.05) was done for the correlation of the primary principal component for anhedonia in the MDD group, using the testnd.Correlation function from the Eelbrain package (50), restricted within the reward > punishment mask. A lower cluster threshold was used due to the smaller sample size for the correlation. Significant clusters are discussed in terms of Brodmann Areas for the sake of interpretation.

## 3. Results

### 3.1 Questionnaires

There were no significant group differences in age, sex, or education, but groups differed on all symptom questionnaires (Table S1). Covariance amongst anhedonia questionnaires was examined using principal components analysis (PCA: Table 1). PCA revealed three distinct factors, with PC1 accounting for anhedonia, PC2 accounting for anxiety and general malaise, and PC3 accounting for apathy. Outcomes were nearly identical when using factor analysis (Table S2).

**Table 1.** Factor loadings from principal components analysis of questionnaires with all subjects (N = 90). DARS and TEPS were reverse coded to represent anhedonia rather than pleasure.

|  | Component |  |  |
| --- | --- | --- | --- |
|  | 1 - Anhedonia | 2 – Anxious Malaise | 3 - Apathy |
| BDI | 0.576 | 0.379 |  |
| BIS |  | 0.583 |  |
| BAS: Drive |  |  | -0.811 |
| BAS: Fun Seeking |  |  | -0.843 |
| BAS: Reward Responsiveness | -0.477 |  | -0.477 |
| SHAPS | 0.699 |  |  |
| DARS: Social | 0.850 |  |  |
| DARS: Hobbies | 0.672 |  |  |
| DARS: Sensory | 0.787 | -0.348 |  |
| DARS: Food | 0.862 |  |  |
| MASQ: Anhedonic Depression | 0.574 | 0.499 |  |
| MASQ: Anxious Arousal |  | 0.810 |  |
| MASQ: General Distress - Depression |  | 0.754 |  |
| MASQ: General Distress - Anxiety |  | 0.908 |  |
| TEPS: Consummatory | 0.622 |  |  |
| TEPS: Anticipatory | 0.676 |  |  |
| AMI: Emotional | 0.429 | -0.401 |  |
| AMI: Social |  |  | 0.644 |
| AMI: Behavioral |  |  | 0.331 |

### 3.2 Behavioral Performance

Healthy controls performed better during the training phase and test phase (Table 2). This learning difference was primarily due to a deficiency in learning from punishments in the MDD group, as assessed by worse NoGo accuracy during the testing phase. There were no significant group differences in PST model parameters or in learning from rewards (Go accuracy). There were no group differences in EEfRT or Pizzagalli task performance (Table S3). Symptom components derived from PCA did not significantly correlate with any task performance measures within the MDD group (Table S4), nor on the EEfRT or Pizzagalli tasks (Table S5). In sum, groups were largely similar on task performance, with the exception of a punishment avoidance deficit in the MDD group.

**Table 2.** Control (CTL) and Major Depression (MDD) participant probabilistic selection task performance and best-fitting model parameters.

|  | CTL | MDD | Statistic and *p* value |
| --- | --- | --- | --- |
| Train: Accuracy (%) | 73.19 (15.443) | 65.06 (13.16) | $t = 2.60, p = 0.011$ |
| Train: Gain learning rate | 0.32 (0.28) | 0.31 (0.32) | $U = 810, p = 0.570$ |
| Train: Loss learning rate | 0.15 (0.28) | 0.21 (0.33) | $U = 874, p = 1.000$ |
| Train: Softmax gain | 4.07 (1.33) | 3.69 (1.80) | $U = 694, p = 0.107$ |
| Test: Accuracy (%) | 70.13 (13.78) | 63.48 (15.89) | $t = 2.03, p = 0.046$ |
| Test: Go accuracy (%) | 66.61 (22.73) | 65.94 (21.36) | $t = 0.139, p = 0.890$ |
| Test: NoGo accuracy (%) | 75.41 (23.38) | 59.65 (28.40) | $t = 2.74, p = 0.008$ |

*Note.* All data are mean ± SD, except model parameters, which are median ± interquartile range. Task performance measures were compared using a Student's *t*-test. Model parameters were compared using a Mann-Whitney U test.

### 3.3 MEG Sensor Analysis

ERFs revealed a RewP analogue from 350-450 ms (Figure 2A). Topographical plots of this temporal region-of-interest (t-ROI) show how frontolateral magnetometers (orthogonal to the midline) had the greatest difference between conditions during this window. The CTL group had a larger reward t-ROI compared to the MDD group (left sensor: t = 2.20, *p* = 0.031; right sensor: t = 2.28, *p* = 0.026; Figure 2B).

**Figure 2.**
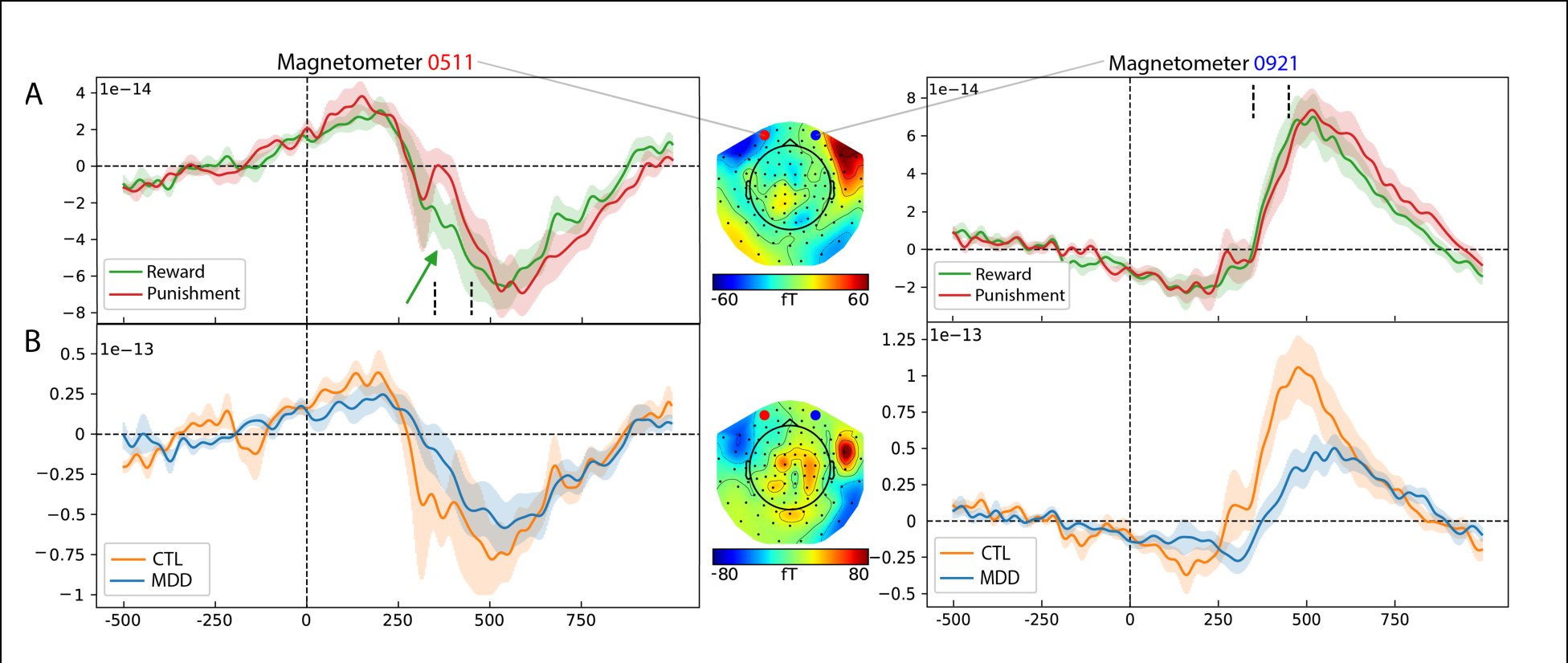
Frontolateral ERFs and topographies A) Feedback locked grand average ERFs following reward and punishment feedback at frontal magnetometers. The topographical plot shows reward > punishment. Sensor “MEG0511” is marked with a red dot and “MEG0921” is marked with a blue dot. **B)** Reward ERFs for each group. The topographical plot shows CTL> MDD.

### 3.4 MEG Source Analysis

Spatiotemporal permutation clustering tests of reward > punishment contrast revealed multiple clusters for reward-specific activation along the medial wall, dorsolateral prefrontal cortex, lateral temporal cortex, and insula (Fig 3). Using these clusters as a spatial mask for reward, independent-sample permutation clustering tests of the orthogonal CTL-MDD contrast revealed a cluster in right ventral cortical areas, encapsulating vmPFC, perigenual cingulate, orbitofrontal cortex, and insula. Time courses of activation for within these major clusters showed similar morphology to the RewP, with greater activation for reward during the t-ROI (Fig S3). Within the MDD group, correlation permutation clustering within this reward mask revealed a significant cluster that was positively correlated with PC1.

**Figure 3.**
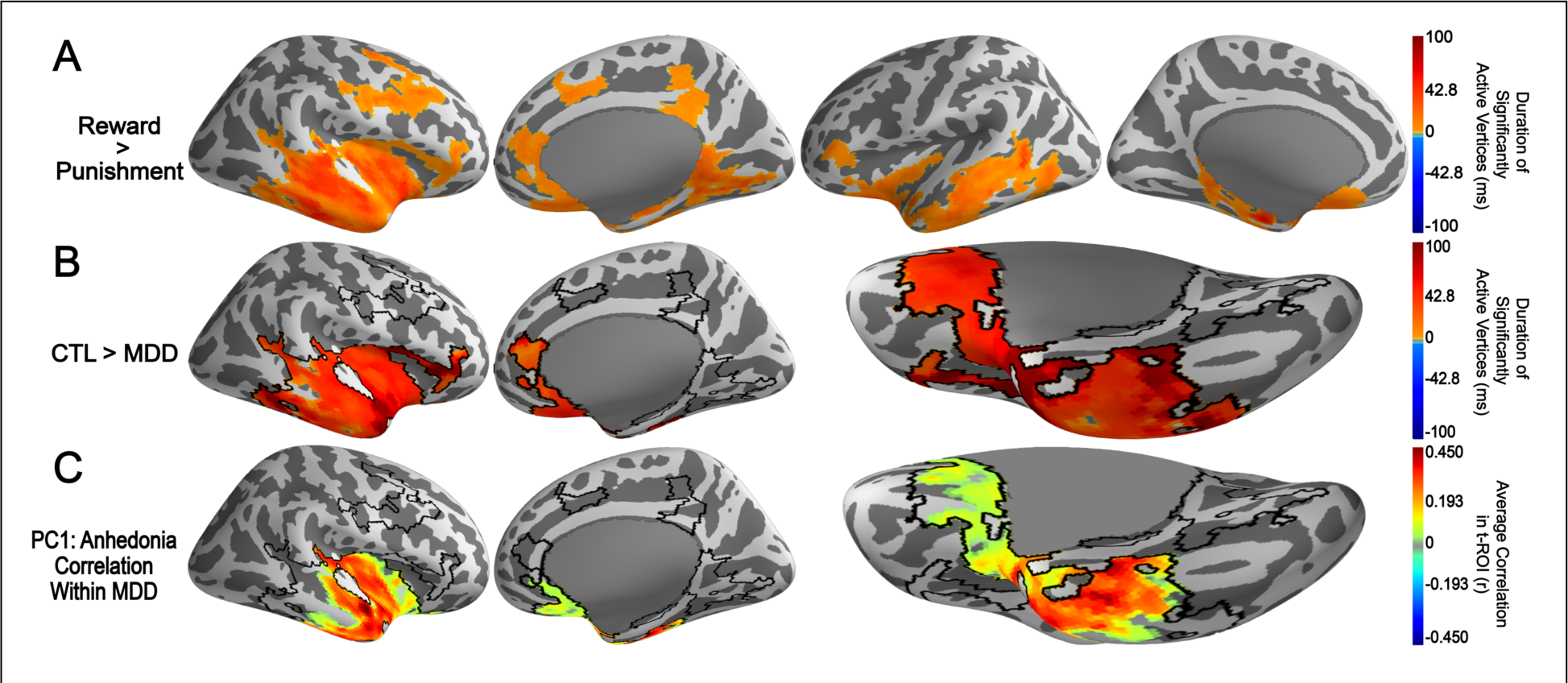
Spatiotemporal permutation clustering test results. A) Significant clusters for reward > punishment feedback in all participants. **B)** Significant cluster for CTL > MDD groups restricted to the reward > punishment mask (black outline). **C)** Significant cluster correlated with PC1 restricted to the reward > punishment mask.

Of the 61 BAs indicated within the reward > punishment clusters, BA25 was the frontal ROI with the largest representation. A majority (85%) of BA25 vertices were in the significant reward > punishment cluster and the CTL > MDD cluster, and 78% of BA 25 was in the PC1 correlation (see Tables S6-S8). Figure 4 shows the time series of this partial BA25 cluster (i.e. the 85% mentioned above), with a clear RewP analogue, a large CTL>MDD difference in the t-ROI (t(79) = 2.69, *p* = 0.009) and a positive correlation with PC1 within the MDD group (*r* = 0.266, *p* = 0.084; smaller than Fig 3C due to collapsing across time in the t-ROI). Correlation of this t-ROI with performance measures revealed an unexpected correlation with aversive learning (NoGo: *r* = 0.302, *p* = 0.049), which was robust across alternative measures and calculations of aversive learning (see Table S9). While both modest and unexpected, we suggest that this behavioral correlation provides important evidence for why anhedonic symptoms are paradoxical to broader MDD group effects.

**Figure 4.**
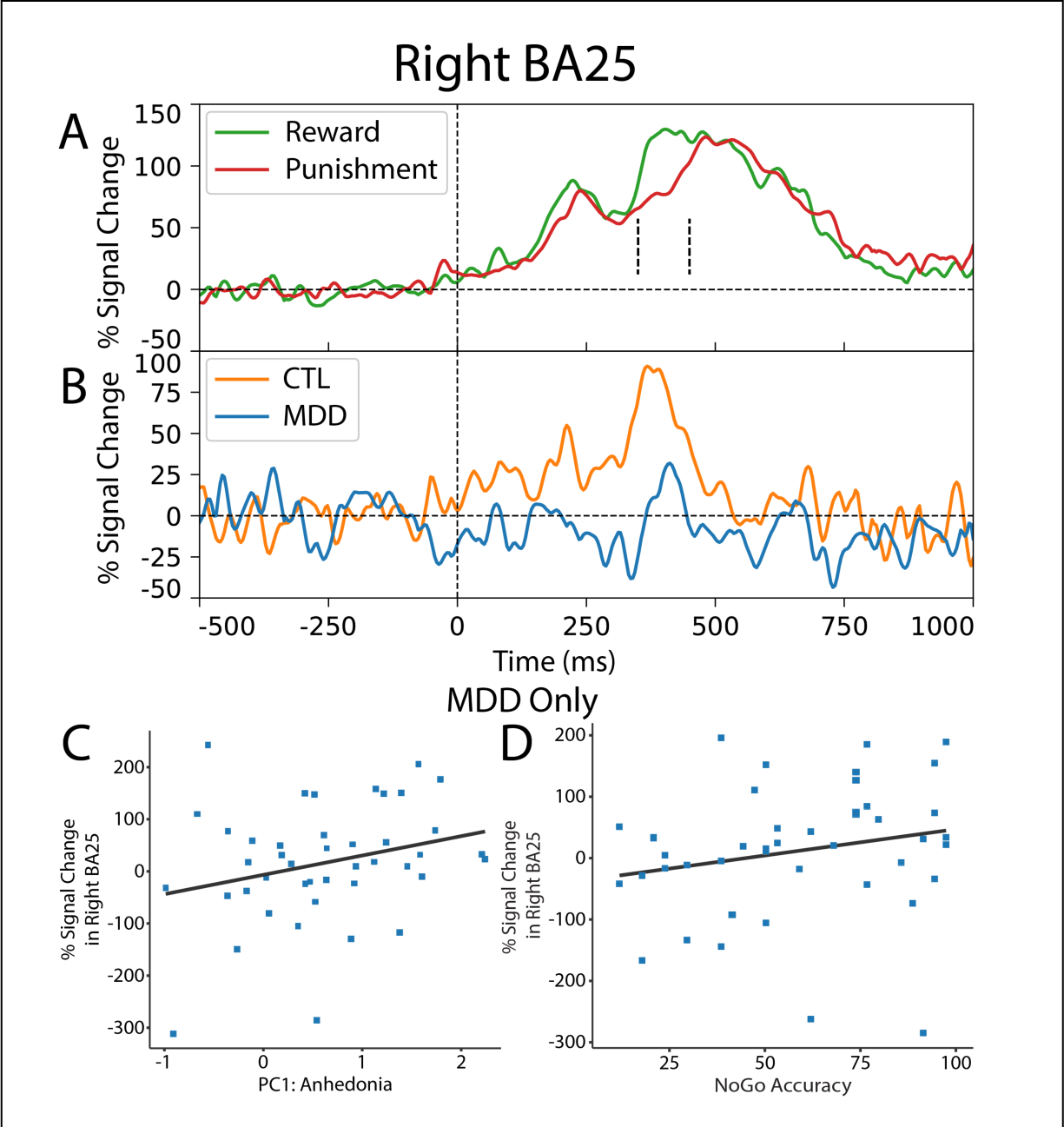
Brodmann Area 25 activation. A) Time course of activation for reward and punishment within partial right BA25 restricted to the reward > punishment mask. **B)** Group differences in the time courses of activation for the reward – punishment difference. In the MDD group, partial right BA25 reward – punishment activity (restricted to significant PC1 cluster) was correlated with both **C)** PC1 anhedonia scores and **D)** NoGo learning. Both of these correlations are paradoxically opposed to the overall group trends, suggesting that anhedonic symptoms reflect a different alteration in reinforcement learning than MDD alone.

## 4. Discussion

This investigation revealed that the cortical response to reward is a product of multiple generative systems, yet mood-related deficits are restricted to ventral areas. This dissociation suggests that different spatial contributors to the RewP provide different information content to the signal. Similar to the extant fMRI literature, this ventral contribution is diminished in MDD, although it is positively correlated with anhedonia. Here we advance a hypothesis about the nature of this paradox, informed by the positive correlation between ventral reward activity and better aversive learning in anhedonia.

Our first major finding was the identification of a robust principal factor of anhedonia across questionnaire measures. This factor was distinct from general malaise and apathy, but it didn’t differ between proposed anhedonic sub-types (e.g. consummatory vs anticipatory). Furthermore, two behavioral assessments of anhedonia failed to scale with this factor or even dissociate groups. These difficulties are not unique to this study, and they demonstrate the challenge of indirect assessments of complex state-relevant affective disturbances. Here we aimed to use a well-known reinforcement learning task to evoke a candidate biomarker of reward to see if it had greater sensitivity to anhedonia.

Our second major finding was that performance on this probabilistic selection task differed between groups, and this was due to altered punishment learning. We previously identified a negativity bias in less severe depressed participants that caused them to over-focus on punishments, leading to *better* punishment learning (51). We later associated that aspect of punishment sensitivity specifically to co-morbid anxiety (23), whereas depressive symptoms were associated with a diminished RewP. In that previous report we proposed that reward-related alterations in depression were not used in an “effective” manner (i.e. to influence learning), but may reflect an “affective” role of reduced hedonic appreciation. We think these predictions were broadly supported here, at least in less severe MDD participants: reward learning was intact even in the context of diminished vmPFC activity. In a moment we will describe how anhedonia adds an additional complexity to this model.

Our third major finding was the first-ever identification of an MEG analogue of the RewP. Fronto-polar sensors showed reward-specific activity in a 90-degree shift from the canonical RewP. These generative sources were distributed across dorsomedial and ventromedial surfaces. Our fourth and most notable major finding was the verification that only ventral sources were different in MDD. The primary frontal contributor to this MDD difference was in BA25: subgenual cingulate. The importance of this region as a target of deep-brain stimulation (52,53) provides convergent evidence that aberrant activity in this area causally underlies the phenotypic expression of depression.

Our final major set of findings identified paradoxical tendencies within the same group: ventral areas were diminished in MDD yet they were positively correlated with anhedonia. Many fMRI studies have found this same pattern (29,31–40), but an explanation has remained elusive Here we propose a testable hypothesis about this anhedonia paradox based on the behavioral correlations discovered here.

### 4.1 Limitations and future directions

We know that in practice, human reinforcement learning includes inevitable cognitive strategies like working memory maintenance and hierarchical goal updating (54,55). Thus, reward receipt is rarely as simple as updating an action value in a “model-free” system. In fact, reward information can be used to update an orthogonal “model-based” goal instead (56,57). Here we showed strong evidence that MDD is generically characterized by diminished reward valuation (lower vmPFC/BA25) as well as compromised punishment avoidance (lower NoGo learning). However, there are simultaneous countervailing tendencies in BA25 as anhedonic symptoms increase. We propose that this could reflect a tendency to shift from a deficient model-free system towards an effective model-based strategy focused on avoidance. Here, the virtue of a reward would be to provide information about the counterfactual (“the other option was the bad choice”). As shown here, the ability to learn from reward may be intact (c.f. Tables 2, S3, S4, S5), but a deficient representation of reward hedonia may instead use reinforcements to update a model characterized by a negativity bias. In other words, rewards offer little more than the absence of punishment. This hypothesis could be tested in a multi-step reinforcement learning task with an anhedonic sample. Importantly, this hypothesis could be successfully tested at low cost using EEG measures of the RewP.

These findings suggest a ventral mood-related deficit in reward receipt, though the spatial scope of this deficit is unclear. Due to the lower number of MEG sensors across ventral areas (the “brim of the cap”), source estimation techniques will spread activity across wide swaths of the cortical depths. This is likely apparent in source estimates spreading from the right ventral midline towards the right ventral temporal lobe. This temporal activation was not predicted by our prior hypotheses, and we interpret it with skeptical caution due to the limits of MEG source estimation. At the limit of this novel method, convergent evidence across multiple imaging assessments will be required to further refine these discoveries. Fortunately, cross-species translational studies offer a new opportunity for understanding this biosignal. Multiple groups have developed rodent analogues of the RewP (58–61). This translational RewP analogue offers a chance to mechanistically test the network-level systems underlying separable reward domains.

### 4.2 Conclusion

The findings reported here add considerable illumination to the nature of an altered RewP in depression. There are multiple distributed source contributions to this signal, and only some of these are affected by MDD. We suggest that the RewP is influenced by separate yet simultaneous cortical computations of reward surprise as well as hedonic liking, identifying it as a nexus where multidimensional value is computed. We further suggest that reward responsivity in anhedonia may differ in complexity from MDD, requiring the assessment of many dimensional symptoms and careful consideration of task design.

The RewP is diminished in depressed individuals (18–22), it can predict the probability of a future major depressive episode in never-depressed individuals (60), and it predicts cognitive behavioral therapy responsivity in depression (63). It has good test-retest reliability (64,65) and internal reliability (66), including reliable diminution due to depressive symptoms (66). These psychometric strengths compliment the aforementioned specificity to reward and sensitivity to multiple aspects of valuation. In sum, the RewP has the sensitivity, specificity, validity, reliability, and prognostic value desired in a biomarker. Here we reported for the first time that BA25, a major dysfunctional node in MDD, is a partial generator of the RewP that shows specific anhedonic alterations, further motivating the RewP analogue as a powerful, prognostic biomarker of mood disturbance.

## Acknowledgements

This project was funded by 1R01MH119382. Research reported in this publication was supported by the Office of The Director, National Institutes of Health of the National Institutes of Health under Award Number S10OD025313. The content is solely the responsibility of the authors and does not necessarily represent the official views of the National Institutes of Health.

## Data Availability

We are currently preparing a BIDS-formatted version of the data and code to upload to Openneuro.org.

## Author Contributions (CRediT)

**CJHP**: Investigation, Project Administration, Data curation, Methodology, Software, MEG Pipeline Development, Formal analysis, Writing – Original Draft

**GS**: Investigation, Project Administration

**JH:** Supervision, Software, Writing – Review & Editing

**DQ**: Supervision, Writing – Review & Editing, Funding acquisition

**JFC**: Conceptualization, Supervision, Project Administration, Methodology, Writing – Original Draft, Funding acquisition

## Supplemental Materials

**Supplemental Figure 1.**
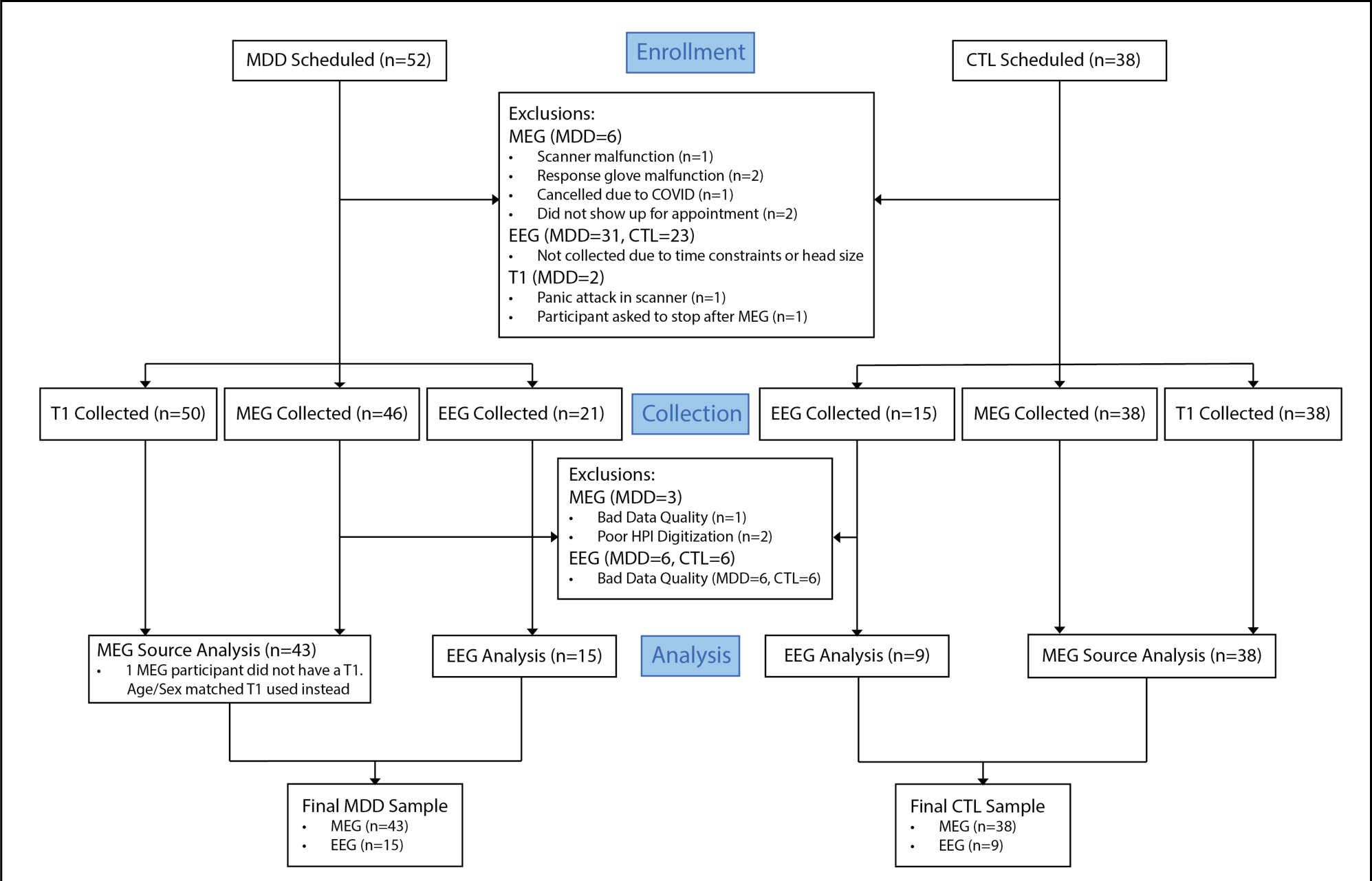
CONSORT Diagram.

**Supplemental Table 1.** Control (CTL: N = 38) and Major Depressive Disorder (MDD: N = 52) participant demographics and average questionnaire scores.

|  | CTL | MDD | Statistic and p value |
| --- | --- | --- | --- |
| Sex | | | $\chi^2 = 1.36, p = 0.244$ |
| Male | 17 | 17 |  |
| Female | 21 | 35 |  |
| Age (years) | 29.68 (3.75) | 27.92 (4.34) | $t = 1.00, p = 0.319$ |
| Education (years) | 15.24 (2.14) | 15.25 (2.76) | $t = -0.02, p = 0.981$ |
| BDI | 4.47 (3.90) | 27.33 (9.78) | $t = -13.62, p < 0.001$ |
| BIS | 20.03 (4.70) | 22.42 (4.06) | $t = -2.59, p = 0.011$ |
| BAS | 42.42 (5.14) | 37.37 (7.12) | $t = 3.73, p < 0.001$ |
| Drive | 11.66 (2.57) | 10.44 (2.66) | $t = 2.17, p = 0.033$ |
| Fun Seeking | 12.32 (2.37) | 10.71 (2.84) | $t = 2.83, p = 0.006$ |
| Reward Responsiveness | 18.45 (1.48) | 16.21 (2.99) | $t = 4.25, p < 0.001$ |
| SHAPS | 0.32 (0.93) | 3.52 (2.59) | $t = -7.29, p < 0.001$ |
| DARS | 60.16 (6.08) | 45.94 (14.22) | $t = 5.72, p < 0.001$ |
| Social | 13.59 (1.80) | 9.75 (4.39) | $t = 5.03, p < 0.001$ |
| Hobbies | 15.43 (0.80) | 11.50 (4.08) | $t = 5.77, p < 0.001$ |
| Sensory | 17.56 (3.15) | 14.43 (5.23) | $t = 3.19, p = 0.002$ |
| Food | 14.05 (1.49) | 11.51 (3.52) | $t = 4.13, p < 0.001$ |
| MASQ |  |  |  |
| Anhedonic Depression | 54.16 (11.63) | 73.60 (14.48) | $t = -6.76, p < 0.001$ |
| Anxious Arousal | 23.76 (6.92) | 32.19 (10.62) | $t = -4.23, p < 0.001$ |
| General Distress – Dep | 21.22 (7.81) | 35.94 (12.16) | $t = -6.47, p < 0.001$ |
| General Distress – Anx | 20.19 (7.09) | 28.02 (8.73) | $t = -4.50, p < 0.001$ |
| TEPS | 83.05 (13.26) | 69.19 (15.15) | $t = 4.51, p < 0.001$ |
| Consummatory | 39.89 (6.84) | 32.90 (7.44) | $t = 4.55, p < 0.001$ |
| Anticipatory | 46.58 (7.96) | 36.83 (11.40) | $t = 4.53, p < 0.001$ |
| AMI | 25.29 (12.34) | 35.38 (9.30) | $t = -4.43, p < 0.001$ |
| Emotional | 7.58 (5.60) | 10.19 (6.72) | $t = -1.95, p = 0.054$ |
| Social | 8.24 (5.00) | 13.04 (4.30) | $t = -4.89, p < 0.001$ |
| Behavioral | 9.47 (5.55) | 12.15 (5.08) | $t = -2.38, p = 0.020$ |

*Note.* All data are mean ± SD. Demographics and questionnaires were compared using a Student's $t$ -test. BDI = Beck Depression Inventory. BIS = Behavioral Inhibition Scale. BAS = Behavioral Activation Scale. SHAPS = Snaith-Hamilton Pleasure Scale. DARS = Dimensional Anhedonia Rating Scale. MASQ = Mood and Anxiety Symptoms Questionnaire. TEPS = Temporal Experience of Pleasure Scale. AMI = Apathy-Motivation Index.

**Supplemental Table 2.**
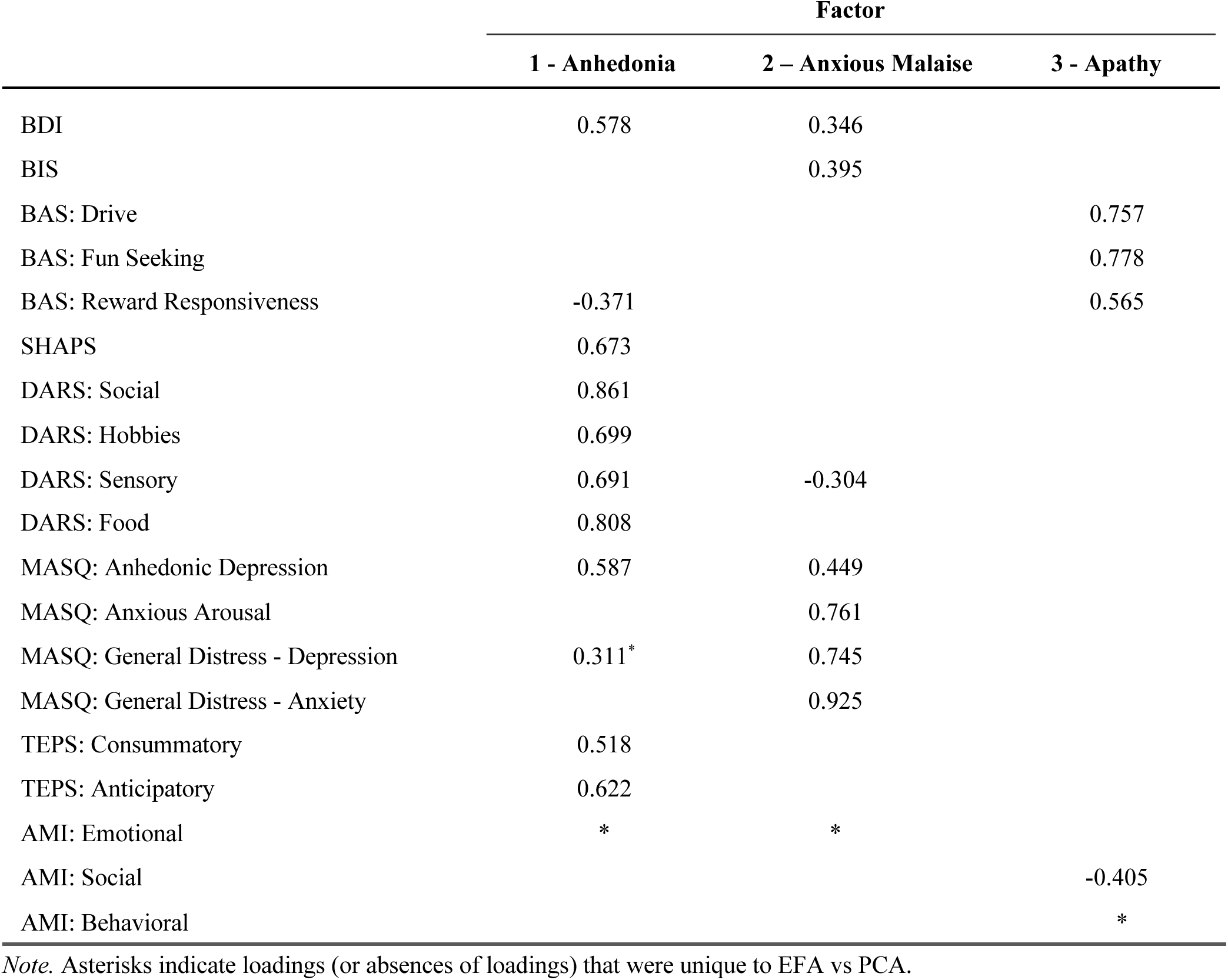
Factor loadings from exploratory factor analysis of questionnaires with all subjects (N = 90)

**Supplemental Table 3.**
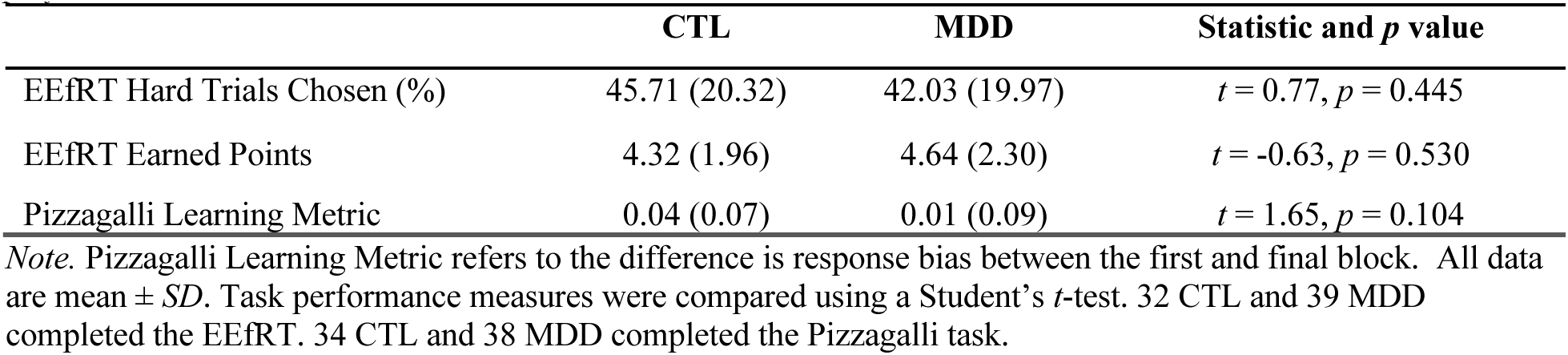
Control (CTL) and Major Depression (MDD) participant EEfRT and Pizzagalli task performance.

| | CTL | MDD | Statistic and $p$ value |
| --- | --- | --- | --- |
| EEfRT Hard Trials Chosen (%) | 45.71 (20.32) | 42.03 (19.97) | $t = 0.77, p = 0.445$ |
| EEfRT Earned Points | 4.32 (1.96) | 4.64 (2.30) | $t = -0.63, p = 0.530$ |
| Pizzagalli Learning Metric | 0.04 (0.07) | 0.01 (0.09) | $t = 1.65, p = 0.104$ |

*Note.* Pizzagalli Learning Metric refers to the difference in response bias between the first and final block. All data are mean ± SD. Task performance measures were compared using a Student's $t$ -test. 32 CTL and 39 MDD completed the EEfRT. 34 CTL and 38 MDD completed the Pizzagalli task.

**Supplemental Figure 2.**
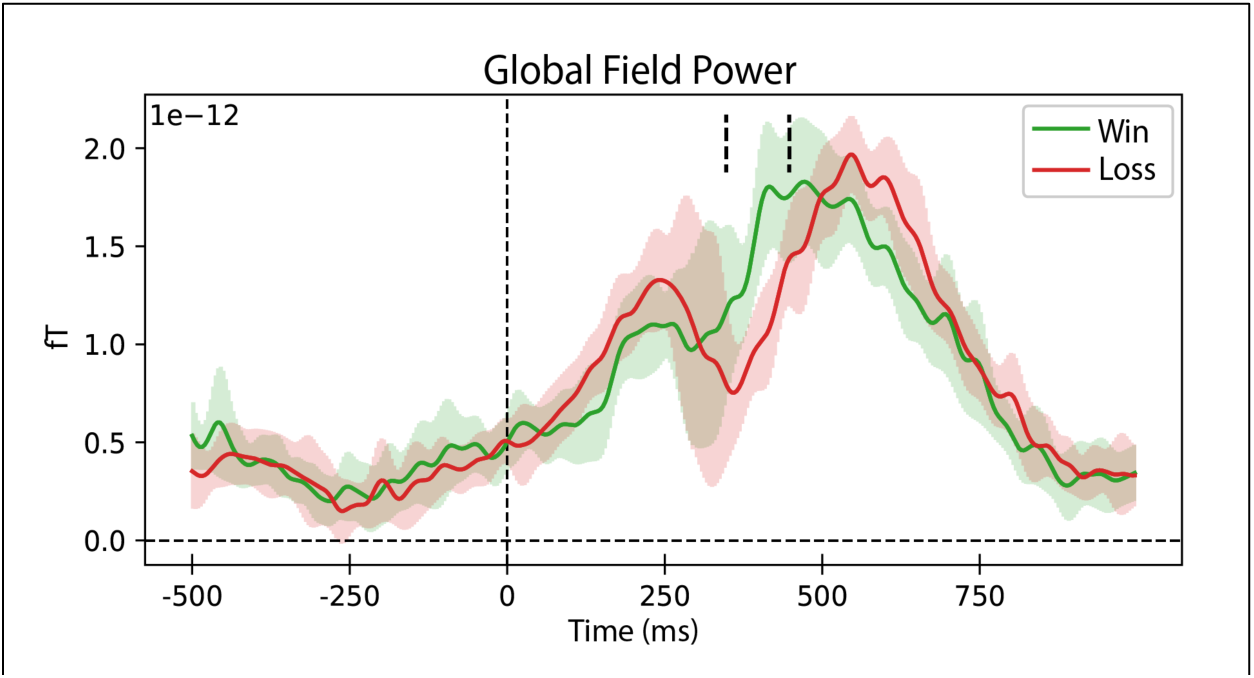
Global Field Power.

**Supplemental Table 4.**
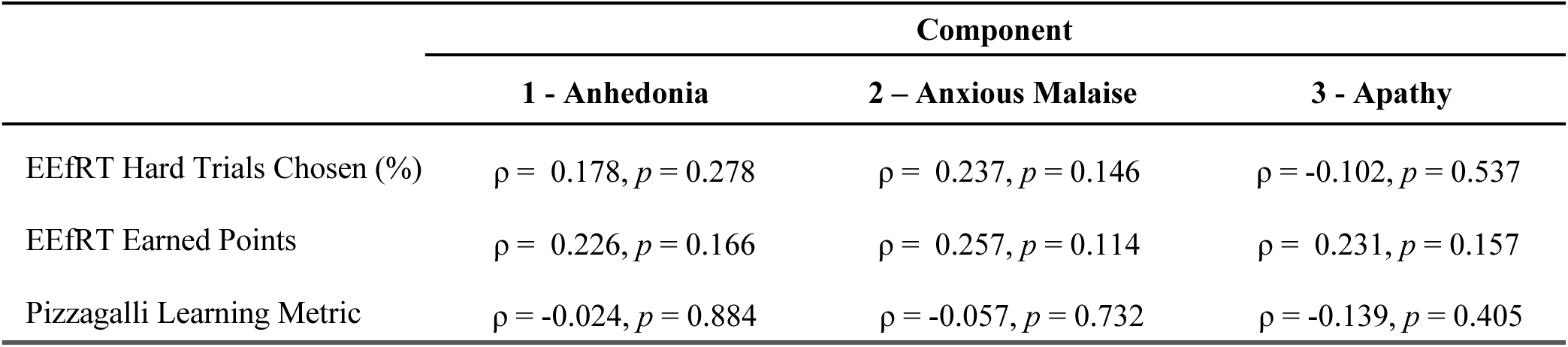
Spearman’s correlation of symptom components with EEfRT (N = 39) and Pizzagalli task (N = 38) performance within MDD subjects.

**Supplemental Table 5.** Spearman’s correlation of symptom components with task performance and best-fitting model parameters in MDD subjects (N = 46)

|  | Component |  |  |
| --- | --- | --- | --- |
|  | 1 - Anhedonia | 2 - Anxious Malaise | 3 - Apathy |
| Train: accuracy | $\rho = 0.037, p = 0.809$ | $\rho = -0.020, p = 0.897$ | $\rho = 0.142, p = 0.348$ |
| Train: Gain learning rate | $\rho = 0.161, p = 0.285$ | $\rho = -0.055, p = 0.714$ | $\rho = 0.168, p = 0.263$ |
| Train: Loss learning rate | $\rho = -0.181, p = 0.227$ | $\rho = -0.200, p = 0.183$ | $\rho = -0.129, p = 0.393$ |
| Train: Softmax gain | $\rho = -0.153, p = 0.308$ | $\rho = -0.034, p = 0.822$ | $\rho = -0.024, p = 0.876$ |
| Test: accuracy | $\rho = -0.046, p = 0.763$ | $\rho = -0.125, p = 0.407$ | $\rho = 0.097, p = 0.520$ |
| Test: A > B accuracy | $\rho = 0.040, p = 0.794$ | $\rho = -0.060, p = 0.694$ | $\rho = 0.126, p = 0.406$ |
| Test: Go accuracy | $\rho = -0.111, p = 0.462$ | $\rho = -0.141, p = 0.350$ | $\rho = -0.017, p = 0.910$ |
| Test: NoGo accuracy | $\rho = 0.074, p = 0.626$ | $\rho = -0.124, p = 0.411$ | $\rho = 0.061, p = 0.687$ |

**Supplemental Figure 3.**
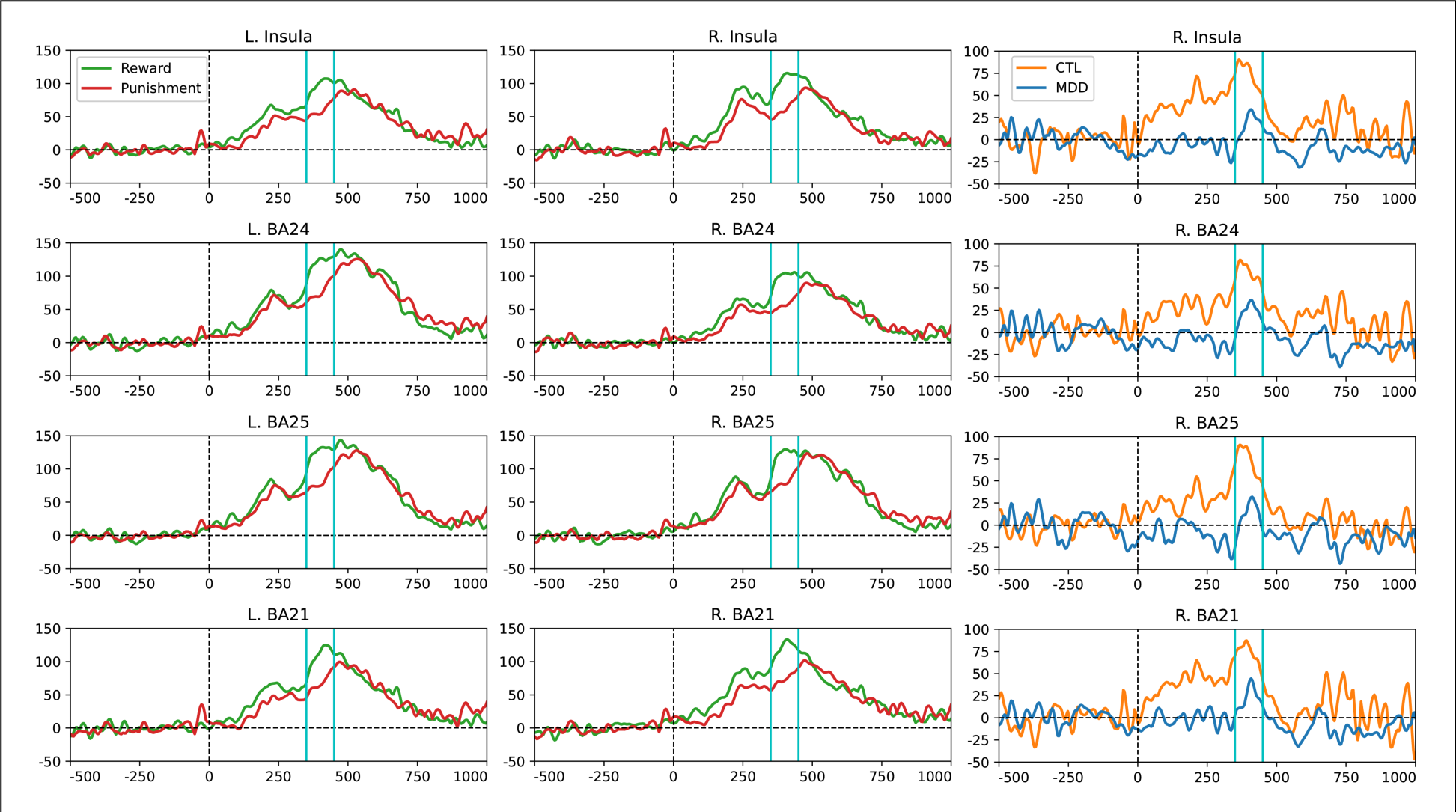
Source-level time courses. Temporal activations for vertices within the significant Reward > Punishment cluster, further grouped within Brodmann Areas. BA21 = Lateral Temporal Cortex; BA24 = Anterior Cingulate Cortex; BA25 = Subgenual Cingulate Cortex. CTL vs. MDD time courses are only shown for right-sided areas.

**Supplemental Table 6.** Reward > Punishment Cluster Overlap with Brodmann Areas.

| ROI | Hemisphere | Region | # Overlapping Vertices | Total ROI Size (Vertices) | % ROI in Cluster |
| --- | --- | --- | --- | --- | --- |
| BA21 | Right | Temporal | 164 | 166 | 99 |
| BA35 | Right | Temporal | 27 | 28 | 96 |
| BA29 | Right | Retrosplenial | 13 | 14 | 93 |
| BA27 | Right | Temporal | 24 | 26 | 92 |
| BA25 | Right | Medial Frontal | 35 | 41 | 85 |
| BA27 | Left | Temporal | 21 | 25 | 84 |
| BA38 | Right | Temporal | 145 | 177 | 82 |
| BA25 | Left | Medial Frontal | 26 | 35 | 74 |
| BA28 | Left | Temporal | 43 | 60 | 72 |
| BA47 | Right | Orbital | 32 | 45 | 71 |
| BA22 | Right | Temporal | 341 | 500 | 68 |
| BA38 | Left | Temporal | 104 | 154 | 68 |
| Insula | Right | Insula | 510 | 789 | 65 |
| BA36 | Left | Temporal | 55 | 86 | 64 |
| BA20 | Right | Temporal | 174 | 276 | 63 |
| BA26 | Right | Retrosplenial | 5 | 8 | 63 |
| BA21 | Left | Temporal | 80 | 137 | 58 |
| BA28 | Right | Temporal | 23 | 41 | 56 |
| BA22 | Left | Temporal | 292 | 524 | 56 |
| BA20 | Left | Temporal | 157 | 297 | 53 |
| BA35 | Left | Temporal | 13 | 25 | 52 |
| BA30 | Right | Retrosplenial | 17 | 40 | 43 |
| BA23 | Right | Posterior Cingulate | 51 | 122 | 42 |
| BA44 | Right | Lateral Frontal | 55 | 133 | 41 |
| BA11 | Left | Orbital | 128 | 311 | 41 |
| BA42 | Right | Temporal | 47 | 120 | 39 |
| BA32 | Right | Medial Frontal | 67 | 175 | 38 |
| BA37 | Right | Temporal | 120 | 335 | 36 |
| BA11 | Right | Orbital | 126 | 354 | 36 |
| Insula | Left | Insula | 242 | 738 | 33 |
| BA45 | Left | Lateral Frontal | 16 | 51 | 31 |
| BA47 | Left | Orbital | 12 | 44 | 27 |
| BA36 | Right | Temporal | 24 | 92 | 26 |
| BA24 | Right | Medial Frontal | 60 | 245 | 24 |
| BA37 | Left | Temporal | 66 | 274 | 24 |
| BA31 | Right | Posterior Cingulate | 48 | 207 | 23 |
| BA6 | Right | Premotor | 115 | 519 | 22 |
| BA46 | Right | Lateral Frontal | 38 | 180 | 21 |
| BA1 | Right | Somatosensory | 33 | 195 | 17 |
| BA3 | Right | Somatosensory | 24 | 154 | 16 |
| BA33 | Right | Medial Frontal | 24 | 154 | 16 |
| BA33 | Left | Medial Frontal | 4 | 26 | 15 |
| BA39 | Left | Parietal | 41 | 281 | 15 |
| BA8 | Right | Frontal | 36 | 253 | 14 |
| BA9 | Right | Frontal | 42 | 299 | 14 |
| BA19 | Right | Occipital | 82 | 637 | 13 |
| BA18 | Right | Occipital | 46 | 386 | 12 |
| BA46 | Left | Lateral Frontal | 21 | 178 | 12 |
| BA39 | Right | Parietal | 29 | 265 | 11 |
| BA29 | Left | Retrosplenial | 1 | 11 | 9 |
| BA32 | Left | Medial Frontal | 15 | 168 | 9 |
| BA4 | Right | Motor | 49 | 566 | 9 |
| BA45 | Right | Lateral Frontal | 6 | 81 | 7 |
| BA17 | Right | V1 | 11 | 195 | 6 |
| BA24 | Left | Medial Frontal | 13 | 263 | 5 |
| BA5 | Right | Parietal | 8 | 168 | 5 |
| BA2 | Right | Somatosensory | 14 | 339 | 4 |
| BA43 | Right | Somatosensory | 2 | 118 | 2 |
| BA40 | Right | Parietal | 3 | 300 | 1 |
| BA40 | Left | Parietal | 3 | 339 | 1 |
| BA19 | Left | Occipital | 3 | 612 | 0 |

**Supplemental Table 7.**
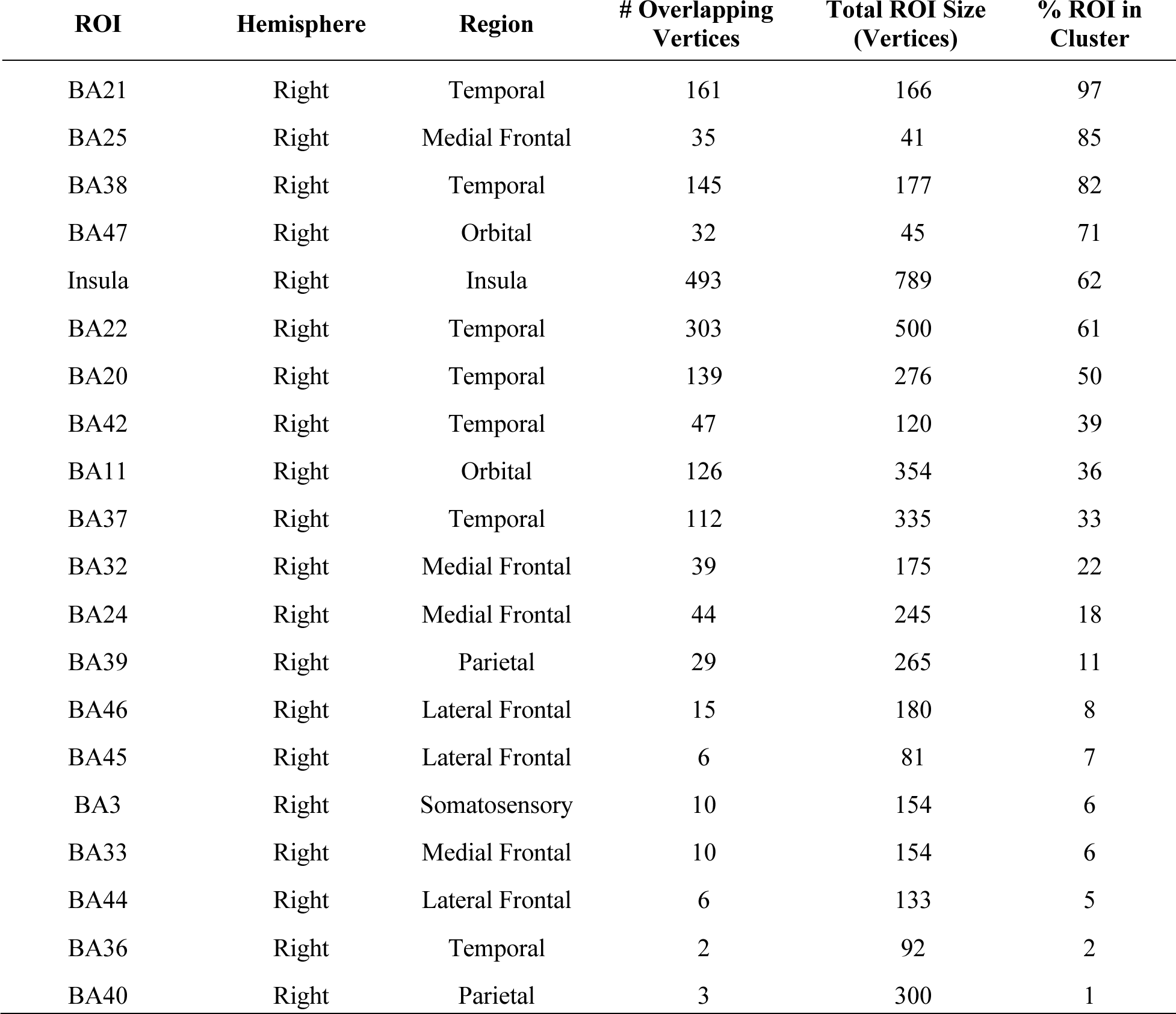
CTL > MDD Cluster Overlap with Brodmann Areas.

| ROI | Hemisphere | Region | # Overlapping Vertices | Total ROI Size (Vertices) | % ROI in Cluster |
| --- | --- | --- | --- | --- | --- |
| BA21 | Right | Temporal | 161 | 166 | 97 |
| BA25 | Right | Medial Frontal | 35 | 41 | 85 |
| BA38 | Right | Temporal | 145 | 177 | 82 |
| BA47 | Right | Orbital | 32 | 45 | 71 |
| Insula | Right | Insula | 493 | 789 | 62 |
| BA22 | Right | Temporal | 303 | 500 | 61 |
| BA20 | Right | Temporal | 139 | 276 | 50 |
| BA42 | Right | Temporal | 47 | 120 | 39 |
| BA11 | Right | Orbital | 126 | 354 | 36 |
| BA37 | Right | Temporal | 112 | 335 | 33 |
| BA32 | Right | Medial Frontal | 39 | 175 | 22 |
| BA24 | Right | Medial Frontal | 44 | 245 | 18 |
| BA39 | Right | Parietal | 29 | 265 | 11 |
| BA46 | Right | Lateral Frontal | 15 | 180 | 8 |
| BA45 | Right | Lateral Frontal | 6 | 81 | 7 |
| BA3 | Right | Somatosensory | 10 | 154 | 6 |
| BA33 | Right | Medial Frontal | 10 | 154 | 6 |
| BA44 | Right | Lateral Frontal | 6 | 133 | 5 |
| BA36 | Right | Temporal | 2 | 92 | 2 |
| BA40 | Right | Parietal | 3 | 300 | 1 |

**Supplemental Table 8.** PC1 Cluster Overlap with Brodmann Areas.

| ROI | Hemisphere | Region | # Overlapping Vertices | Total ROI Size (Vertices) | % ROI in Cluster |
| --- | --- | --- | --- | --- | --- |
| BA38 | Right | Temporal | 143 | 177 | 81 |
| BA25 | Right | Medial Frontal | 32 | 41 | 78 |
| BA21 | Right | Temporal | 121 | 166 | 73 |
| BA20 | Right | Temporal | 139 | 276 | 50 |
| Insula | Right | Insula | 388 | 789 | 49 |
| BA42 | Right | Temporal | 47 | 120 | 39 |
| BA22 | Right | Temporal | 141 | 500 | 28 |
| BA11 | Right | Orbital | 70 | 354 | 20 |
| BA37 | Right | Temporal | 42 | 335 | 13 |
| BA24 | Right | Medial Frontal | 18 | 245 | 7 |
| BA32 | Right | Medial Frontal | 10 | 175 | 6 |
| BA3 | Right | Somatosensory | 4 | 154 | 3 |
| BA33 | Right | Medial Frontal | 4 | 154 | 3 |
| BA36 | Right | Temporal | 2 | 92 | 2 |

**Supplemental Table 9.**
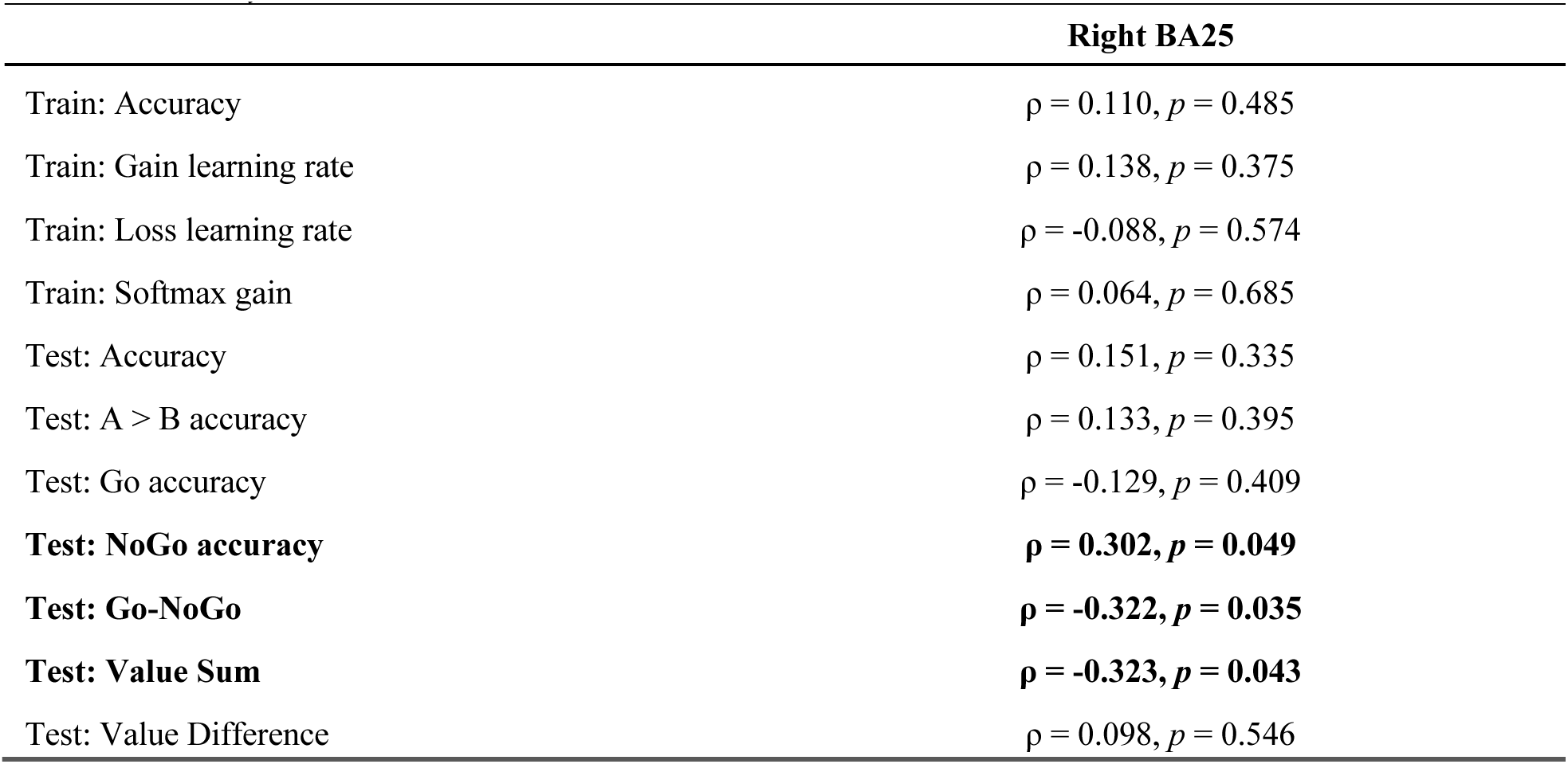
Spearman’s correlation of partial Right BA25 (Reward > Punishment) activity with task performance and best-fitting model parameters in MDD subjects (N = 43). Alternative measures of test phase selection are based on logistic beta weights for Value sum (added values of the two stimuli) and Value difference (subtracted values of the two stimuli). Value difference is highly correlated with overall accuracy whereas Value sum is more closely related to the Go-NoGo measure.

